# Enhancing tree seed germination prediction with image-driven machine learning models

**DOI:** 10.1101/2025.01.21.634084

**Authors:** Pablo Gómez Barreiro, Adam Richard-Bollans, Efisio Mattana, Charlotte E. Seal, Ted Chapman, Roberta Dayrell

**Affiliations:** Royal Botanic Gardens, Kew, Wakehurst, Ardingly, Haywards Heath, West Sussex RH17 6TN, UK; Royal Botanic Gardens, Kew, Richmond, London, TW9 3AE, UK

**Keywords:** UK native species, forestry, machine learning, deep learning, seed germination, computer vision, seed morphology, ecological restoration, trait variability

## Abstract

1. Tree planting is crucial for reversing deforestation and meeting net zero targets, requiring a reliable supply of high-quality seeds. Efficient use of limited native seeds can be promoted by sorting methods, but traditional techniques commonly used for agricultural species are often unsuitable for tree seeds due to their high trait variability.
2. Here, we explored the potential of combining image analysis with machine learning models to improve tree seed sorting outcomes. We selected five UK native tree species of interest for tree production and afforestation projects and applied machine learning XGBoost and Convolutional Neural Networks algorithms to predict seed germination using colour and X-ray images as well as features extracted from these images.
3. The machine learning models achieved good accuracy and F1-scores, but their specificity was limited, particularly when relying solely on colour images or related features. This poses a problem, as wild seeds are often scarce, and falsely classifying seeds that germinate as non-germinable would result in a waste of valuable resources. X-ray images and features were highly effective in identifying empty seeds but did not perform well when differentiating filled seeds into germinable and non-viable. Consequently, the models performed best for species with a high proportion of empty seeds.
4. For three of the five species, model performance varied significantly by mother tree, with some trees showing markedly poorer results. This aspect had not been previously investigated and raises concerns that biased seed sorting will disadvantage certain mother trees, leading to the loss of valuable genetic diversity and woodland resilience.
*Synthesis and applications*: The performance of image-based machine learning models in predicting seed germination ultimately depended on whether most non-germinated seeds were empty, non-viable, or dormant. X-ray models showed strong performance in detecting empty seeds, but colour image models exhibited poor results due to the high variability in seed external features, the subtle differences between germinated and non-germinated seeds, and the variability among individual mother trees. Developing open, accessible training databases and more adaptable models is crucial for addressing these limitations and enable technologies to further support large-scale tree production.

## Introduction

Nearly one-third of the Earth’s land is covered by forests, of which 7 percent (290 million hectares) is planted (FAO, 2020). Tree planting plays a fundamental role in reversing global net deforestation trends and meeting net zero targets, but current restoration efforts demand a vast supply of native seeds (Merritt & Dixon, 2011; Nevill et al., 2018). To meet these ambitious targets (UN DESA, 2019), it is of paramount importance to ensure a reliable source of high quality and genetically diverse seeds is available (Jalonen et al., 2018), a role significantly played by the seed industry.

Seed sorting methods are commonly used to enhance crop seed quality and germination predictability, resulting in more productive supply chains. Large-scale commercial differentiation between "good" (germinable and highly vigorous) and "bad" (non-viable, empty or infested) seeds for agricultural crops is accomplished using non-destructive methods such as sieving, gravity tables, and optical sorting. Bred for consistency, lack of dormancy, and ease of cultivation, crop seed traits are of homogeneous nature, facilitating the detection of non-germinable (e.g. immature, insect damaged or malformed) and low-quality seed outliers by using a range of morphological traits as benchmarks (Rahman & Cho, 2016, and references therein). On the other hand, seeds from non-domesticated native trees show much greater variability in seed traits (including seed dormancy) due to the influence of greater genetic diversity and variability in environmental conditions during seed production on the mother plant (Callejas-Díaz et al., 2022; Cendán et al., 2013; Goszka & Snell, 2020). Therefore, sorting methods cannot be directly applied to native tree seeds without undesirable outcomes, such as discarding viable seeds, which are a scarce and valuable resource, and artificially selecting seeds in ways that may reduce genetic diversity. These methods must be adapted to effectively address the unique characteristics of non-domesticated populations, where maintaining trait and genetic diversity is essential to guarantee the adaptive potential and resilience of the populations.

The recent surge in solutions that integrate image analysis with machine learning (ML) for crop seed sorting (Kumar et al., 2024) has introduced new possibilities for overcoming the limitations of traditional optical seed-sorting methods for wild tree species. ML models can assess multiple seed traits simultaneously, capturing subtle differences that traditional, rule-based sorting systems (often limited to simpler traits like size or weight) might miss (Pichler & Hartig, 2023). By learning from a diverse range of features such as colour, texture, and shape, ML models can offer a more flexible approach than manual methods, enabling the analysis of tree seed traits and their relationships to germination potential (Rahman & Cho, 2016).

However, if not handled correctly, this variability could also introduce bias into seed sorting, potentially favouring seeds from specific trees or with certain morphological traits, thereby reducing genetic diversity in seed batches used in restoration (Hesami et al., 2022; Schaberg et al., 2008). The extent of this potential bias in ML models is still unclear and represents a crucial gap in the field that requires further investigation.

In addition to sorting challenges, seeds from wild species also tend to fail to germinate more frequently than those of agricultural crops, due to factors such as the absence of embryos, non-viable embryos, or dormancy mechanisms, each presenting distinct morphological characteristics. X-ray and colour imaging offer complementary insights to help distinguish different classes of non-viable seeds from potentially germinable ones. X-ray images provide a non-invasive view of internal seed structures, allowing the detection of seeds lacking embryos or containing damaged tissues, which are good indicators of seed viability, especially for seeds with little or no endosperm (Tausch et al., 2024). In contrast, colour images capture external traits such as shape, size, texture, and colour, and can also indicate seed quality, for example, by allowing the identification of mouldy seeds (Sun et al., 2016). Viable seeds that are dormant or lose viability during germination trials are difficult to differentiate based on visual traits alone, as they are viable but may not survive and germinate under certain conditions. The ability of these imaging methods to reliably distinguish between seeds able to germinate from non-viable and empty seeds has yet to be fully explored, particularly for tree species (Bianchini et al., 2021). Whether integrating these two imaging methods can enhance seed sorting efficiency also remains an open question.

This study seeks to explore the potential of combining colour and X-ray imaging with ML models to handle the high variability in wild seed traits and reliably predict germination. We selected five species of interest for UK tree production and use in afforestation projects, and aimed to answer the following questions: Can image-driven ML models effectively distinguish seeds that will germinate from non-viable and empty seeds? How effective are X-ray images when integrated with ML? Can external seed traits, such as size and colour, reliably predict wild seed viability? Additionally, we examine whether ML models exhibit biases based on individual mother trees, potentially affecting genetic diversity in seed batches used for afforestation and restoration. By tackling these questions, this study aims to uncover both successes and limitations of image-based seed sorting technologies, offering insights that could pave the way for improved practices to meet the increasing demand for high-quality, genetically diverse tree seeds essential for global restoration efforts.

## Material and methods

### Tree seed material

We selected species with contrasting morphological features, along with two species of similar morphology, to enable comparisons and generalised conclusions on the application of image-based ML methods across different taxa. Seeds of *Alnus glutinosa* (L.) Gaertn, *Betula pendula* Roth, *Betula pubescens* Ehrh., *Pinus sylvestris* L. and *Sorbus aucuparia* L. were collected across the UK between 2016 and 2018, with the seeds from each individual mother tree kept separate (Chapman et al., 2019). Seeds were dried at 18°C and 15% relative humidity, cleaned and placed in dry and cold storage (15% relative humidity and -20°C) at the Millennium Seed Bank (Royal Botanic Gardens, Kew) until the start of the experiments in 2021. Samples consisted of approximately 1,000 seeds for each species, with the number of seeds taken from each mother tree proportional to the original collection, thereby preserving the relative contribution of seeds from each mother tree to the overall sample (Table 1).

**Table 1.**
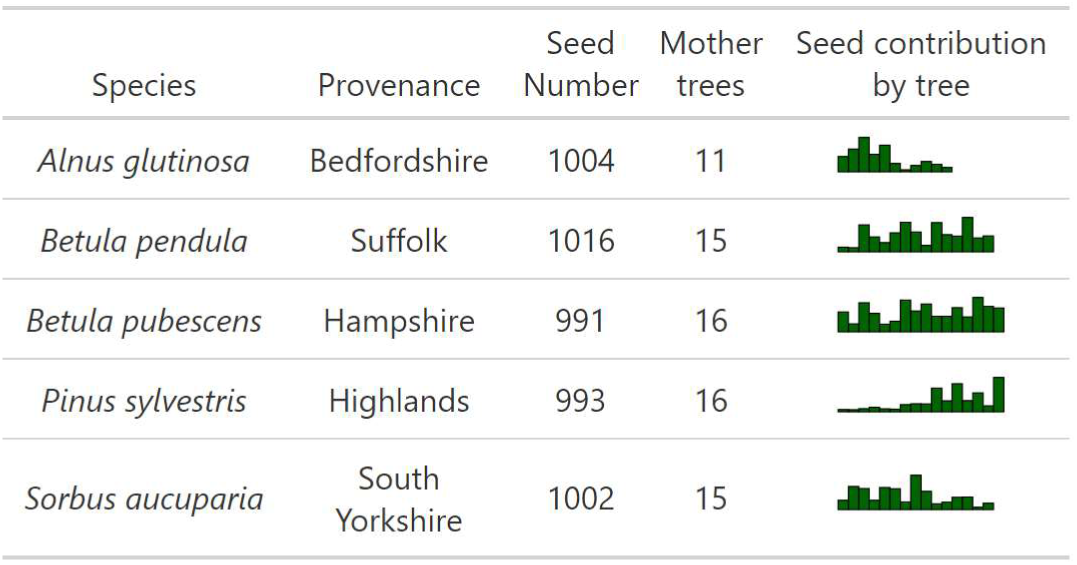
Summary of used seed material: Provenance, total number of seeds analysed (Seed N), number of mother trees sampled, and contribution to the population by mother tree.

| Species | Provenance | Seed Number | Mother trees | Seed contribution by tree |
| --- | --- | --- | --- | --- |
| *Alnus glutinosa* | Bedfordshire | 1004 | 11 |  |
| *Betula pendula* | Suffolk | 1016 | 15 |  |
| *Betula pubescens* | Hampshire | 991 | 16 |  |
| *Pinus sylvestris* | Highlands | 993 | 16 |  |
| *Sorbus aucuparia* | South Yorkshire | 1002 | 15 |  |

### Morphological characterization

Each seed was placed in a fixed position within a 9x9 grid. This arrangement allowed for consistent tracking of each seed, keeping information on the mother tree from which it was collected as well as data from all assessments: X-ray imaging, photographing, germination, and cut tests (Fig. 1).

**Figure 1.**
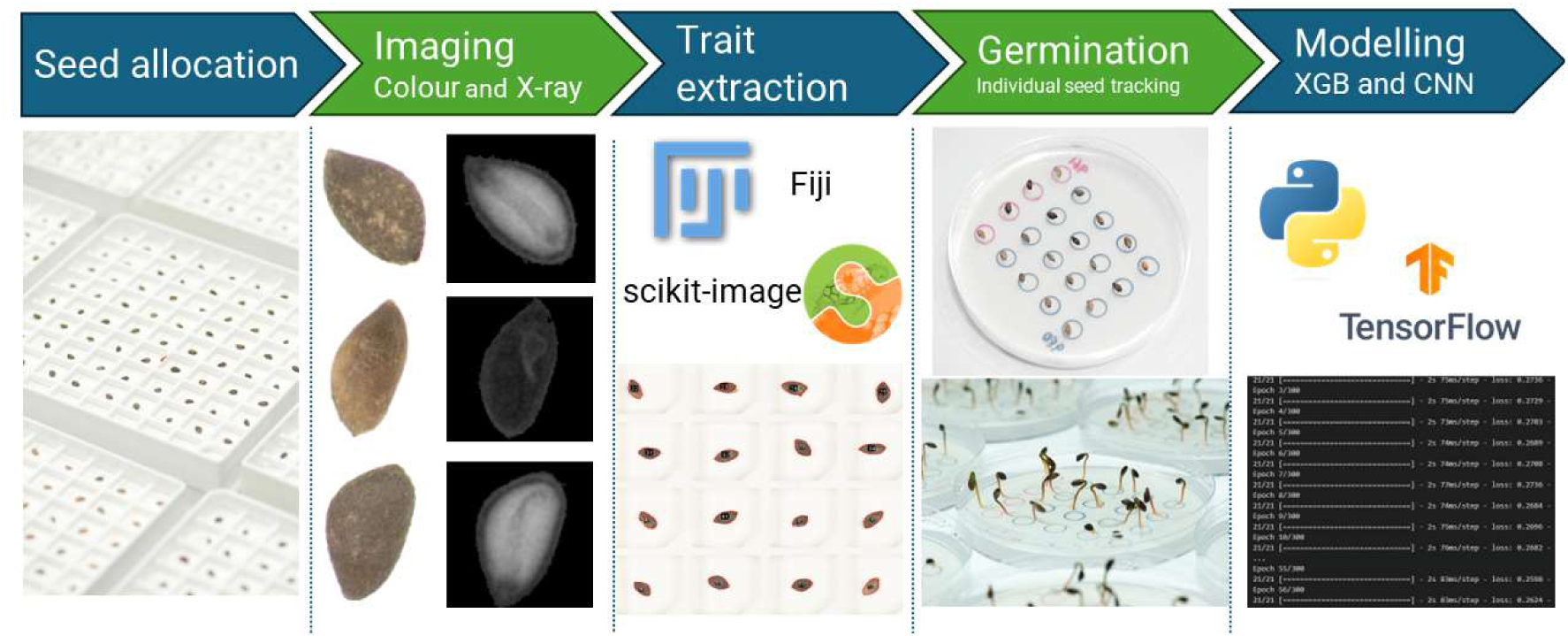
Summary of methods used to obtain seed morphological data for subsequent training.

Grids were photographed with X-rays (Ultrafocus, Faxitron; settings detailed in Table S1a) and standard photography (105mm Nikkor lens mounted on a Nikon D750 body; settings detailed in Table S1b) attached to a copy stand with an illuminated base and fluorescent top lights (eVision executive HF, Kaiser Fototechnik). All images were processed using Fiji (Schindelin et al., 2012). Seeds were identified as ‘regions of interest’ within each image and individually segmented from the background to determine morphometric (area, perimeter, maximum and minimum ferret diameter, circularity, roundness, aspect ratio and solidity), and colour traits (L, a and b channels from CIELAB scale, and average and core grey colour from X-ray images).

Seed texture features (contrast, dissimilarity, homogeneity, energy, angular second moment and correlation) were obtained from converting individual images to grayscale with 32 levels (Haralick et al., 1973). A grey-level co-occurrence matrix (GLCM) considering invariant spatial direction (as an average of all four angles: 0°, 45°, 90°, 135°) was then constructed to capture the spatial relationships of pixel intensities. This matrix was normalised and used to calculate features, which were computed using Scikit-image (Van Der Walt et al., 2014) library functions, graycoprops and graycomatrix.

### Seed germination

After imaging and morphological characterisation, each seed was transferred to a preallocated position within a petri dish (each containing 30 ml. 1% water/agar and 25 seeds), allowing the germination status of each individual seed to be tracked and linked to morphological traits (Fig. 1). We incubated seeds under conditions conducive to germination based on Davies et al. (2020) and data from Millennium Seed Bank’s Database, summarized in Table S2. Germination was scored every day until experiment completion for *A. glutinosa*, *B. pendula*, *B. pubescens* and *P. sylvestris*, while ***S. aucuparia*** germination was checked on a weekly basis. Germination experiments were concluded two weeks after the last observed germination. For model training and evaluation purposes, seeds were classified into two groups: "0" for seeds that did not germinate and "1" for seeds that germinated. Additionally, the remaining seeds were dissected and visually inspected and classified as fresh (potentially viable), mouldy (non-viable), or empty.

### Statistical analysis

The parametric distribution of morphological traits was tested using the Shapiro-Wilk normality test (Shapiro & Wilk, 1965). Due to non-normal data distribution, pairwise comparisons were performed with the Mann-Whitney U test (Mann & Whitney, 1947), and p-values were adjusted using the Holm correction (Holm, 1979).

### Machine Learning models

XGBoost (Chen & Guestrin, 2016) and Convolutional Neural Network (CNN) algorithms were trained on the binary classification of seed germination outcomes (“1”, germination, “0”, no germination). To determine the most informative dataset in predicting germination, we trained XGBoost models with the morphological features (detailed in the "Morphological Characterization" section) extracted from photographs, X-ray images and the combination of both. Hereafter, these models are referred as XGB colour, XGB X-ray and XGB all, respectively. Each model was trained and evaluated on data from each species individually, as well as on data from all species in the study combined. In each case, a hold-out set was generated for final model evaluation using an 80-20 stratified split into train-test data. Hyperparameters of the XGBoost models were tuned via cross-validation on the training set using GridSearchCV, implemented in the scikit-learn Python library (Pedregosa et al., 2011), with 5 stratified folds using a range of parameters (Table S3). The best hyperparameters were selected based on F1 scores and the models were retrained on the entire training set with these parameters prior to predictions.

The CNN model for image classification was built using TensorFlow (Martín Abadi et al., 2015). Colour and X-ray individual seed images were split 80-20 using the same stratification as for XGBoost. Unlike XGBoost, the models were trained exclusively with either color or X-ray images. To configure the model and associated hyperparameters (early stopping, learning rate, number of training epochs, layer types etc..), the training data was further divided into training and validation data using an 80-20 split. Internal validation performance of the models was iteratively improved by inspecting the training and validation loss, F1 score, accuracy and precision as well as the epoch vs. loss curves and ROC curves. The final model included the following layers: augmentation, rescaling, three convolutional layers with 16, 32, and 64 filters respectively, each followed by a MaxPooling2D layer. These were followed by a dropout layer with a rate of 0.2, after which the output was flattened and passed through a dense layer with 128 units and ReLU activation, culminating in a final dense layer with a single unit and sigmoid activation for binary classification. The model was trained using binary crossentropy loss and the RMSprop optimizer for a maximum of 500 epochs with a learning rate of 0.0001, and an early stopping depending on validation loss.

To assess how effectively these models classify seeds into germinated (accept) and non-germinated (discard) categories, and how much they can enhance sorting methods, we calculated the final germination percentage of the accept subset and examined the condition of seeds in both resulting groups. The final predictive performance of the models was formally evaluated on the test set based on accuracy, F1 score and specificity. To assess whether the model performance would vary according to mother trees, these metrics were also obtained for each tree.

### Software

Data analysis and visualization were conducted using Python (v. 3.11) and R (v. 4.4.1, R Core Team, 2024). The latter included the following packages: ***ggfx*** v.1.0.1 (Pedersen, 2022), ***ggh4x*** v.0.2.8 (Brand, 2024), ***ggtext*** v.0.1.2 (Wilke & Wiernik, 2022), ***gt*** v.0.11.1 (Iannone et al., 2024), ***hrbrthemes*** v.0.8.7 (Rudis, 2024), ***patchwork*** v.1.3.0 (Pedersen, 2024), and ***tidyverse v.2.0.0*** (Wickham et al., 2019).

## Results

### Seed germination and composition of non-germinated fractions

The proportion of germinated seeds varied among the five species studied, ranging from 93% of the total number of seeds for *S. aucuparia* to 59% for *B. pubescens* (Table S4, and Fig. 2). The composition of the remaining non-germinated seeds also varied from species to species. Most seeds did not germinate due to being empty in *A. glutinosa*, whereas *B. pubescens*, *B. pendula*, and *P. sylvestris* non-germinated seeds had a more balanced distribution between empty and mouldy seeds. Only a small fraction of non-germinated but potentially viable seeds was present in *S. aucuparia* (3.6%), *B. pubescens* (3%), and *P. sylvestris* (1%), indicating that the dormancy-breaking protocols and germination conditions used were effective in promoting the germination of viable seeds.

**Figure 2.**
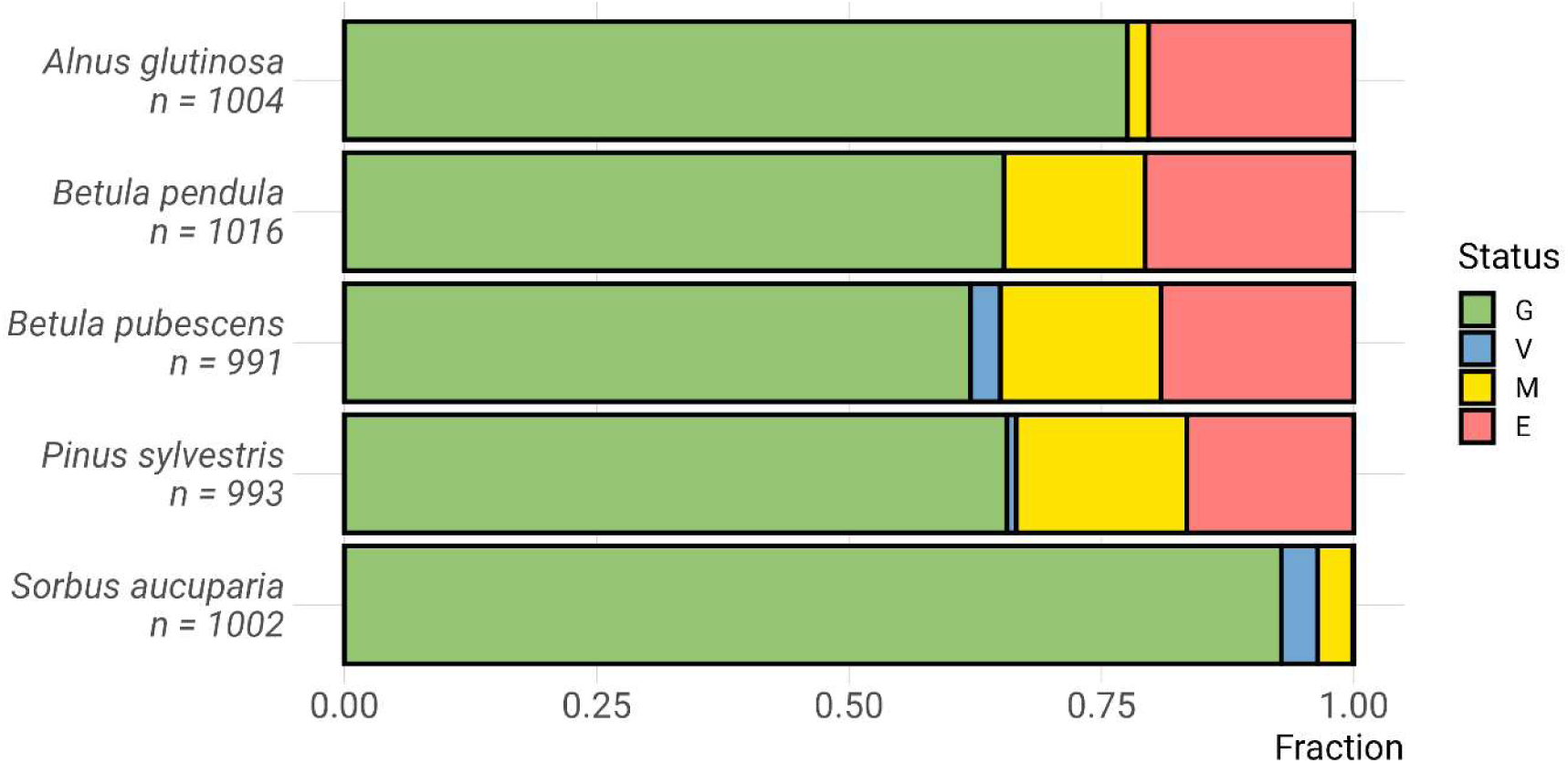
Fractions of germinated (G), non-germinated but viable (V), mouldy (M) and empty (E) seeds at the end of the germination experiments for each of the species studied. n = number of seeds for each species.

### Model-enhanced seed collection quality

As shown in figure 3, X-ray-based single-species models provided effective classification of seeds into germinated (accept) and non-germinated (discard) categories for all species except *S. aucuparia*, thereby enhancing final germination of the accept subsets compared with unsorted seeds. *A. glutinosa* had the largest increase in final germination (from 75% to 97%), with its discard subset consisting mainly of empty seeds. X-ray models also successfully removed empty seeds from *P. sylvestris, B. pubescens, and B. pendula*, leaving the accept subset free of such seeds in the former species. However, these models were not efficient at removing mouldy seeds in any of the species, as most of them remained in the accept subset. X-ray models combining all species were effective at eliminating empty seeds from the accept subset but, similar to models built for single species, struggled to correctly classify mouldy seeds as discard.

**Figure 3.**
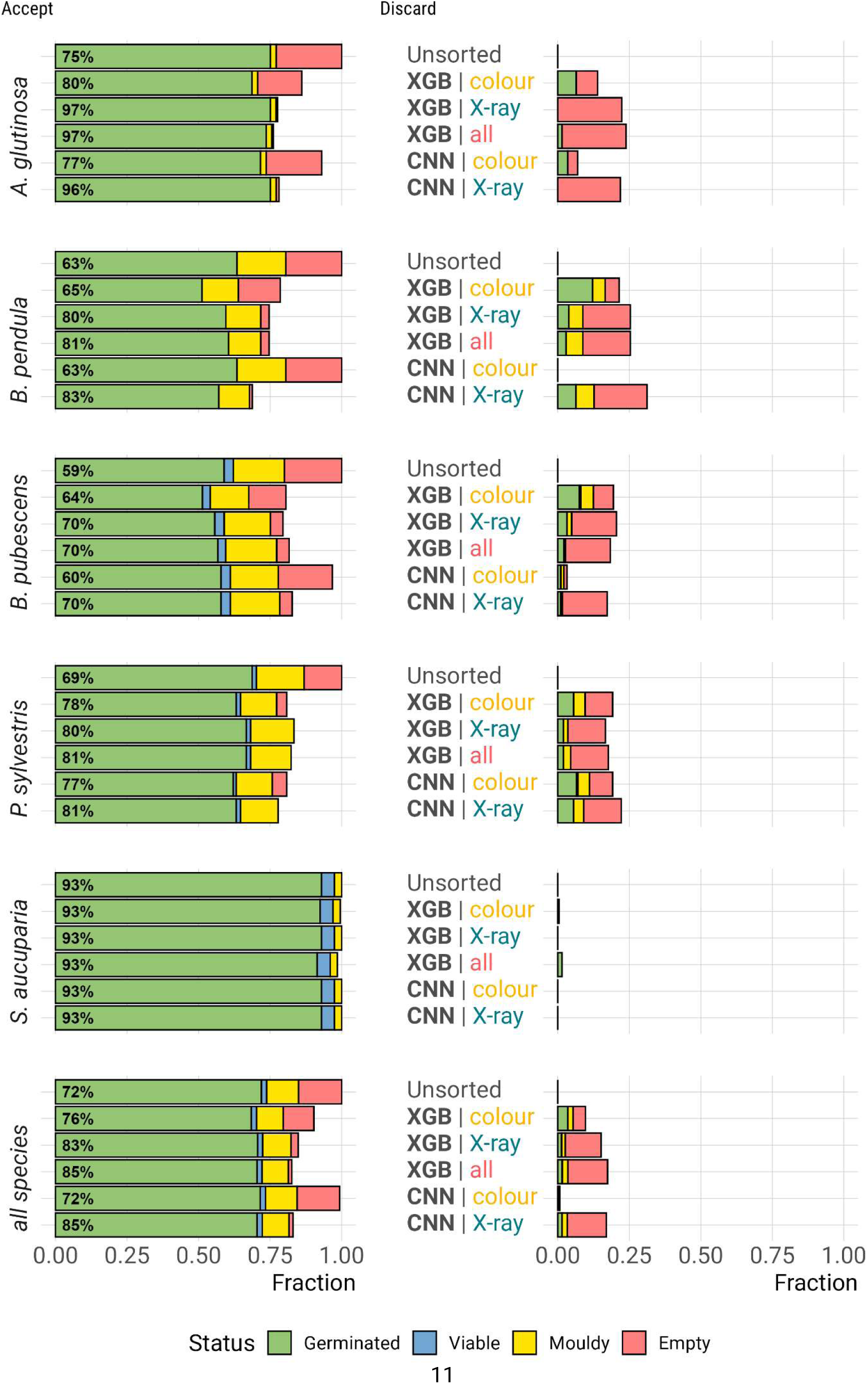
Predicted effectiveness of image-based models in sorting seeds into two subsets: ’Accept’ (left), predicted to germinate, and ’Discard’ (right), predicted not to germinate. All non-germinated seeds were classified as viable, non-viable, or empty. X-axis values indicate the fraction relative to the total number of seeds. Left: Initial status of seeds used for model evaluation (Unsorted) and predicted status after applying different models. Percentages within green bars represent final germination rates within unsorted or accept subset. Right: Predicted status of discarded seeds after the application of each model.

Combining colour and X-ray images for XGBoost was generally as effective as models that used only X-ray images. Additionally, models relying exclusively on colour images performed considerably worse than their X-ray-based counterparts. Overall, colour-only models were less effective at assigning non-germinated seeds to the discard subset and were more prone to misclassifying germinated seeds within it. In certain cases, such as with CNNs for *B. pendula* and all species combined, the models failed entirely to predict any seeds as non-germinated, resulting in the absence of a discard subset. All seeds in the discard subsets of *S. aucuparia* were germinated, regardless of the model.

The evaluation metrics for the trained XGB and CNN models indicated better performance when X-ray data were included compared to using only colour images (Fig. 4). Among all X-ray based models, whether using a single species or a combination of all, *A. glutinosa* achieved the highest accuracy, F1, and specificity scores (≥0.9). The remaining species also had robust accuracy (> 0.73) and F1 scores (>0.81), and specificity values ranging from 0.39 to 0.68. The only exception was *S. aucuparia*, which had specificity scores of zero across all models. Models trained on datasets containing X-ray information from all species combined also showed robust scores for evaluation metrics, but overall, these scores remained around the average values observed for individual species.

**Figure 4.**
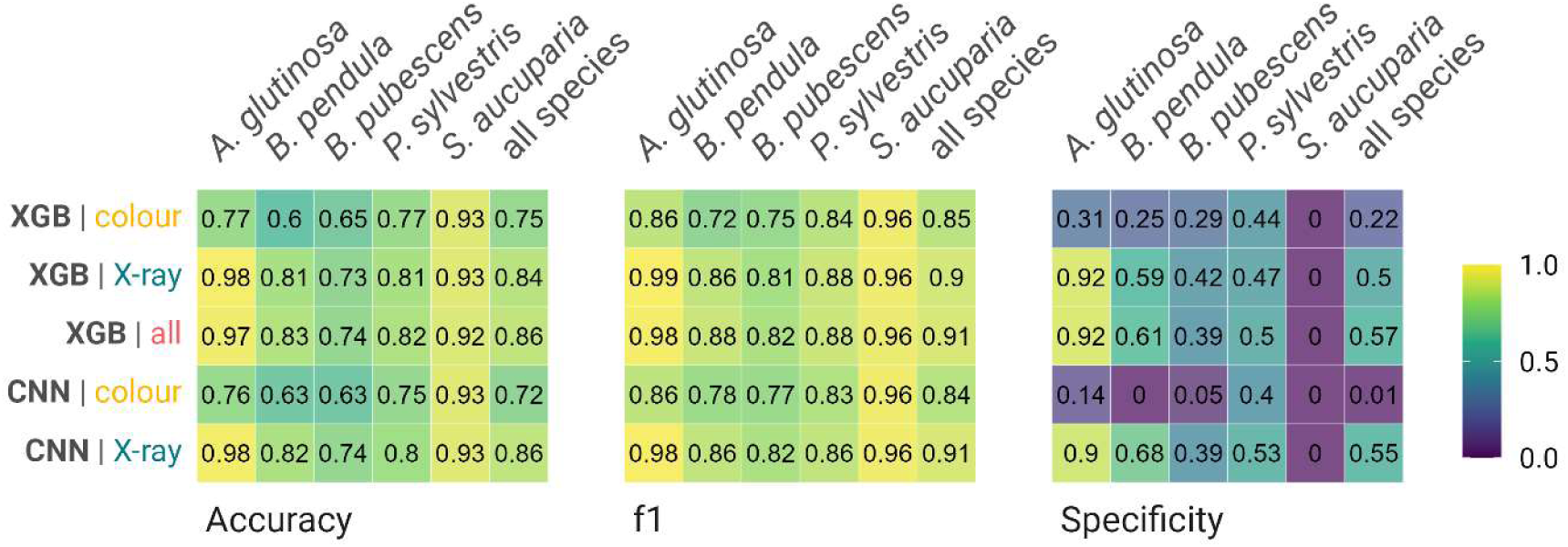
Summary of metrics (Accuracy, f1 and Specificity) collected from evaluation of trained models (XGB, CNN) on testing dataset using traits obtained from colour and Xray images and the combination of these traits (all).

### Traits and metric for species

We found significant differences in the distribution of seed morphological trait values between germinated and non-germinated seeds across the five species (Fig. S1). Germinated and non-germinated seeds differed in 17 traits for ***A.*** *glutinosa*, 13 traits for *P. sylvestris*, 12 traits for *S. aucuparia*, 9 traits for *B. pubescens*, and 8 traits for *B. pendula*. Only the morphological traits obtained from the CIELAB colour channel a (*Mean_a*) in the colour images, and the average grey value of the seed centroid (*Mean_core_grey*) in X-ray images were consistently significantly different across all species. Despite these significant differences, the magnitude of differences is small, and trait distributions exhibit considerable overlap between groups for most traits, with notable separation observed only in the traits obtained from X-ray images (*Mean_grey* and *Mean_core_grey*) for some species.

### Traits and metrics for mother trees

Including the mother tree information in the trait analysis did not enhance the separation between germinated and non-germinated seeds beyond what was observed at the species level. Once again, notable differences between groups were found only for the *Mean_grey* and *Mean_core_grey*, while the other traits continued to exhibit considerable overlap (Fig. S2).

Moreover, the proportions of germinated to non-germinated seeds differed between mother trees, with some extreme cases such as *B. pendula* tree 10 and *P. sylvestris* tree 16 (Fig. S2). Both had a higher proportion of non-germinated seeds in comparison with other trees in the population.

Model performance at the level of individual mother trees varied according to species and training datasets (Fig. S3). There was little variation in performance among mother trees for *A. glutinosa* X-ray-based models, and all *S. aucuparia* models. However, for all other species and for the *A. glutinosa* datasets with only colour images, performance varied considerably by mother tree, with some trees exhibiting markedly lower scores than others in the population. There was not a clear relationship between model metrics for individual mother trees and the proportion of seeds of the population included in the training dataset.

## Discussion

In this study, we assessed the effectiveness of machine learning models using XGBoost and CNN architectures to predict germination of UK native tree species based on colour and X-ray images, and features extracted from these images. While the models showed good accuracies and F1-scores, their performances were limited by low specificities, especially in those relying solely on colour images and their features. This poses a problem, as wild seeds are often scarce, and a sorting method that frequently misclassifies seeds that germinate as non-germinable will result in a substantial waste of valuable resources. On the other hand, X-ray-based models proved to be highly effective for classifying empty seeds as non-germinable, but less so for differentiating between germinated and non-viable (mouldy) seeds. Consequently, the models performed exceptionally well with ***A. glutinosa*** across all metrics, as most of its non-germinated seeds were empty, with only a small proportion being filled but non-viable. A similar outcome is observed in other studies, where the non-germinating fraction primarily consists of empty seeds (De Medeiros et al., 2021; Hamdy et al., 2024). Thus, assessing whether non-germinated seeds are empty or non-viable is essential for understanding the model’s performance and determining the relevance of the method for each species. This insight can also help guidie the selection of seed sorting methods to be tested in future research. Lastly, we showed that for three of the five species, model performance varied significantly by mother tree, with some trees in the population performing markedly worse, an aspect of this type of sorting approach that had not yet been investigated. This could lead to biased seed sorting that may disadvantage mother trees with poorer performance, potentially causing the loss of valuable genetic diversity (Schaberg et al., 2008).

As anticipated, models using internal seed structures as input outperformed those relying only on external morphology. The latter contributed little to the combined models, offering no or minimal additional value. Although some external traits obtained from colour images showed significant differences between germinated and non-germinated seeds, there was considerable overlap in values for most traits. This inherent variability and the subtle distinctions between seeds that can germinate and those that cannot made these traits poor predictors for the models in general. This lack of differentiation may even be linked to a protective strategy, where the similarity between potentially viable and empty seeds helps reduce predation pressure (Myczko et al., 2015; Perea et al., 2013).

Despite external morphological data of tree seeds not significantly enhancing our models, future research should still investigate the potential of image-derived morphological data, such as those captured across various wavelengths (Feng et al., 2019; Pang et al., 2021) and through imaging of surface topography (Martinez et al., 2021), in combination with ML. Unlike crop seeds, the use of hyperspectral imaging and ML for quality sorting of tree seeds remains largely unexplored, despite some research indicating promising potential (Bianchini et al., 2021). Moreover, crop seeds benefit from extensive and diverse image datasets that facilitate the development of more powerful and adaptable models, such as convolutional neural networks combined with transfer learning techniques (Nehoshtan et al., 2021). In contrast, our research on tree seeds is hindered by the absence of large-scale databases, limiting our ability to employ these robust models. This disparity in data availability highlights the urgent need to build comprehensive, shared databases for tree seeds, which could unlock the potential for more advanced algorithms. Researchers should prioritize making raw images and associated data publicly available to foster further advancements in the field.

While X-ray trained models effectively classified empty seeds as not able to germinate, they were unable to accurately distinguish between filled seeds that germinated and those that did not. For ***B. pendula***, ***B. pubescens***, and ***P. sylvestris***, a considerable proportion of non-viable seeds were misclassified as germinated and different factors may contribute to this. Non-dormant seeds without visible damages may fail to germinate due to genetic degradation, accumulation of damage during maturation, lack of antioxidants, or mechanical injuries (Long et al., 2015). Moreover, even when the damage is visible, the extracted features may fail to capture this information (Silva et al., 2012). As the extent and location of damage can vary considerably, resulting in highly variable appearances of non-viable seeds, a larger image dataset is needed for model training to improve their accurate detection. An important aspect of this finding is that studies on tree seeds and less-or non-domesticated species that rely solely on visual inspection of embryos for assessing germination potential (e.g. Dumont et al., 2015) may overestimate germination outcomes and thus, misclassify a significant proportion of seeds with no germination potential. For the small fraction of potentially healthy seeds classified as fresh after the cut test, improvements in the stratification treatments might have ensured their germination. However, it cannot be ruled out that these seemingly viable seeds, like the others, could have become non-viable if the experiments were extended.

Models combining data from all species did not outperform individual species models, as their metrics tended to reflect an average across all species. While multi-species models may capture broader trends, they often fail to optimize for species-specific traits, likely due to interspecies variability introducing noise or confounding patterns. Expanding the dataset could, however, help models generalize seed quality patterns more effectively while minimizing the impact of species-specific variability.

Our study shows that both model types perform similarly when trained on X-ray data. XGBoost models are more cost-effective, requiring less computational power and offering greater interpretability. While feature extraction adds an extra step to the XGBoost models, extending the pipeline, this can be optimised through automated approaches (Dayrell et al., 2023). CNNs, by contrast, automatically extract features from images, reducing manual input but at a higher computational cost and with lower interpretability (Jiang & Li, 2020). Thus, both models perform similarly in seed sorting, but their application by the industry would ultimately depend on available resources and user preferences.

Finally, our work investigated seed sorting performance based on individual mother trees, an aspect that has been largely unexplored. Maintaining adequate levels of genetic variability is a core objective of restoration projects (Di Sacco et al., 2021), but it is subject to seed availability in an overstrained supply chain (Merritt & Dixon, 2011; Nevill et al., 2018). Seed suppliers play a critical role in guaranteeing an adequate supply of genetically diverse seeds by making efforts to source from different trees within populations. However, these efforts can be undermined during seed cleaning processes. In this study, we showed that even a state-of-the-art technology for sorting, such as image-based ML approaches, misclassified a large proportion of the germinated seeds of particular trees in the population as non-germinated. In practice, this would have largely reduced the planting of seeds from those trees. Thus, such practices may inadvertently exclude germinable seeds with uncommon morphological traits, potentially altering gene frequencies and reducing the representation of rare alleles (Schaberg et al. 2008). These alleles are essential for population adaptation and resilience to environmental change. Therefore, we argue that this is a matter that deserves further attention in future studies and highlight the need to develop new approaches that prioritize the preservation of genetic diversity in trees, mitigating the risks associated with seed sorting practices both in wild populations and seed orchards.

Our study highlights the potential of image-based machine learning models to improve seed sorting, thereby reducing resource wastage and increasing cost-effectiveness of tree production and restoration. Our findings show that X-ray models are highly effective for detecting empty seeds across all species and are particularly suitable for collections where non-germinated seeds are predominantly empty. With *A. glutinosa*, a collection meeting this condition, the method was predicted to increase final germination rates from 75% in the original collection to over 95% in the sorted subset, with minimal misclassification of viable seeds. Additionally, we encountered several challenges that are likely common across non-domesticated species. These include frequent misclassification of non-viable seeds and the poor performance of colour-based models across all tested species, both of which stem from the inherent variability in seed traits and the limited availability of datasets. Advanced techniques like transfer learning, while powerful, require large, annotated image sets that are resource-and time-intensive to create, and this is especially challenging for wild species due to much fewer economic incentives. This highlights the urgent need for collaborative efforts to build open, accessible databases to enhance seed sorting practices of wild species globally. Moreover, future research should explore new imaging and other high-throughput methods to identify better traits for distinguishing seed viability. At the same time, seed sorting practices must balance improving germination rates with preserving genetic diversity, which is critical to ensure evolutionary potential. By advancing phenotyping methods and creating shared resources, we can develop scalable, robust models that not only ensure seed quality but also support ecosystem resilience in large-scale restoration projects.

## Supporting information

FigS1_high_resolution

FigS2_high_resolution

FigS3_high_resolution

## Author contributions

**Conceptualization:** PGB, ARB, RD. **Data curation:** PGB, ARB, RD. **Formal analysis:** PGB, ARB, RD. **Investigation:** PGB, ARB, RD. **Methodology:** PGB, ARB, RD. **Software:** PGB, ARB, RD. **Writing (original draft, review and editing):** PGB, ARB, EM, CS, TD, RD.

## Acknowledgements

The authors would like to express their appreciation for the support provided at various stages of this study by our colleagues Owen Blake, Vicky Philpott, Rachael Davies, David Coleshill, and Kaitalin White.

## Data availability

Data, images and code utilized for the training of XGBoost and CNN models, and the analysis of their outputs are available in Figshare (DOI: 10.6084/m9.figshare.28247336) and GitHub (https://github.com/pgomba/ml_tree_seeds).

---

**Table S1a.** Summary of used manual X-ray parameters.

| Species | Exposure | Contrast thresholds |
| --- | --- | --- |
| *Alnus glutinosa* | 5.8 sec. at 23Kv | 4506 to 6295 |
| *Betula pendula* | 6.6 sec. at 20Kv | 4151 to 3233 |
| *Betula pubescens* | 6.6 sec. at 20Kv | 4151 to 3233 |
| *Pinus sylvestris* | 6.2 sec. at 26Kv | 5951 to 9657 |
| *Sorbus aucuparia* | 5.8 sec. at 23Kv | 4506 to 6295 |
| All images were originally saved in .tiff format |  |  |

**Table S1b.** Summary of manual parameters used in standard photography. Constant parameters: Distance from camera to grid: 78 cm, Copystand top light: On.

| Species | Settings | Copystand base light intensity |
| --- | --- | --- |
| *Alnus glutinosa* | 1/50, f9, ISO200 | 3.5 |
| *Betula pendula* | 1/50, f9, ISO200 | 3.5 |
| *Betula pubescens* | 1/50, f9, ISO200 | 3.5 |
| *Pinus sylvestris* | 1/50, f9, ISO250 | Off |
| *Sorbus aucuparia* | 1/50, f9, ISO200 | 3.5 |
| Prior to image analysis, original NEF photographs were converted to .tiff |  |  |

**Table S2.** Summary of germination conditions.

| Species | Stratification | Germination Temp. |
| --- | --- | --- |
| *Alnus glutinosa* | 5°C – 41 days | 25°C |
| *Betula pendula* | 5°C – 39 days | 25°C |
| *Betula pubescens* | 5°C – 41 days | 25°C |
| *Pinus sylvestris* | No | 20°C |
| *Sorbus aucuparia* | No | 5°C |
| Sources: Davies et al., 2020 and MSB's Seed Bank Database |  |  |

**Table S3.** XGBoost grid search hyperparameters used in the internal loop of the nested cross validation and the final model.

| Hyperparameters | Values |
| --- | --- |
| learning_rate | 0.01, 0.05, 0.1, 0.3 |
| n_estimators | 100, 500, 1000 |
| subsample | 0.3, 0.5, 0.9 ,1 |
| min_child_weight | 1, 5, 9, 11 |
| gamma | 0, 1, 4 |
| colsample_bytree | 0.6, 0.8, 1 |
| max_depth | 0.6, 0.8, 1 |

**Table S4.** Germinated and non-germinated seed composition used to train and evaluate the different models.

| Species | Set | Germinated |  |
| --- | --- | --- | --- |
|  |  | Yes | No |
| *Alnus glutinosa* | Train | 630 | 173 |
|  | Test | 152 | 49 |
| *Betula pubescens* | Train | 534 | 277 |
|  | Test | 130 | 75 |
| *Betula pendula* | Train | 492 | 300 |
|  | Test | 123 | 76 |
| *Pinus sylvestris* | Train | 516 | 279 |
|  | Test | 136 | 62 |
| *Sorbus aucuparia* | Train | 743 | 58 |
|  | Test | 187 | 14 |
| all species (Total) | Train | 2915 | 1087 |
|  | Test | 728 | 276 |

**Figure S1.**
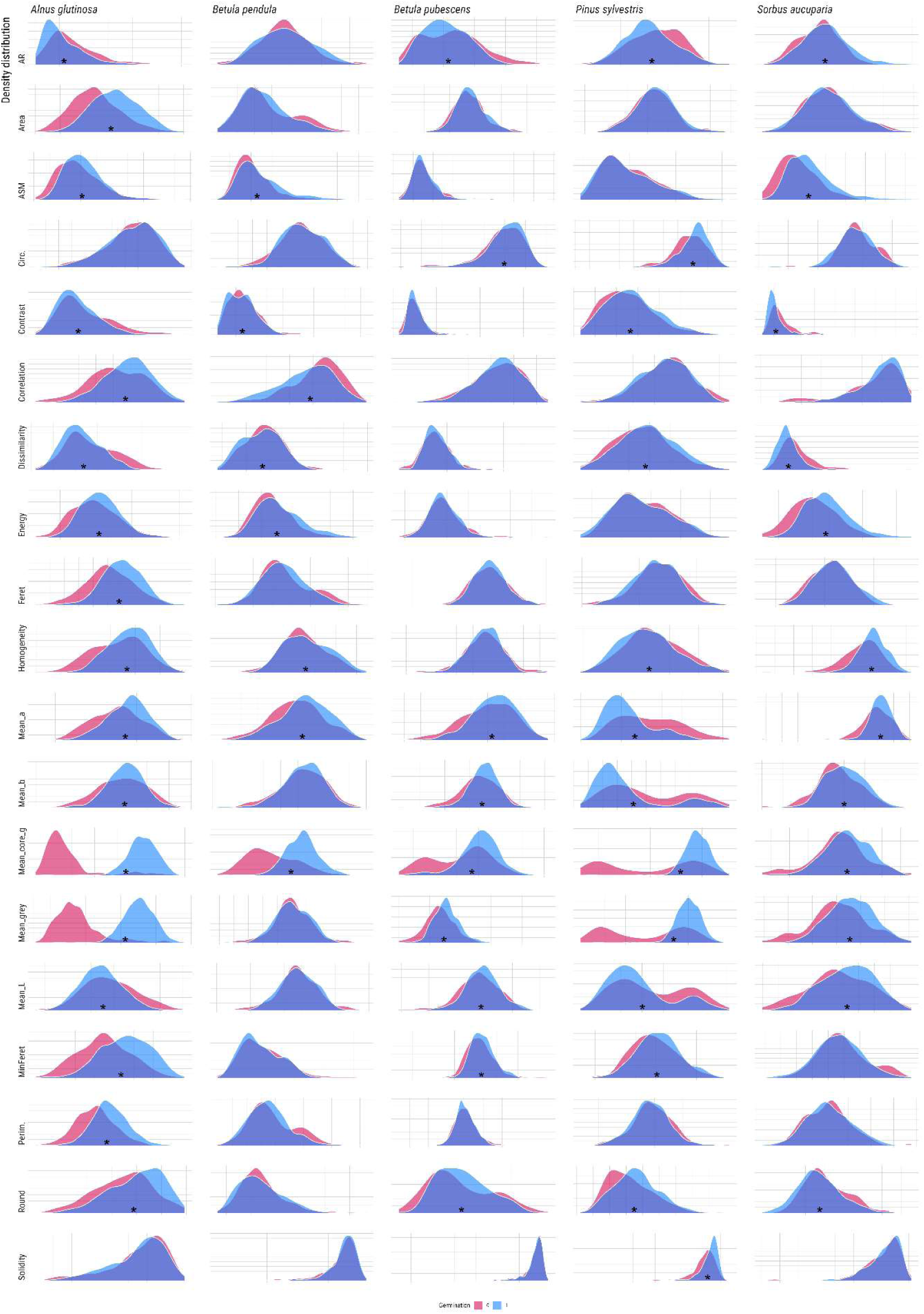
Distribution of seed morphological trait values between germinated and non-germinated seeds across all species

**Figure S2.**
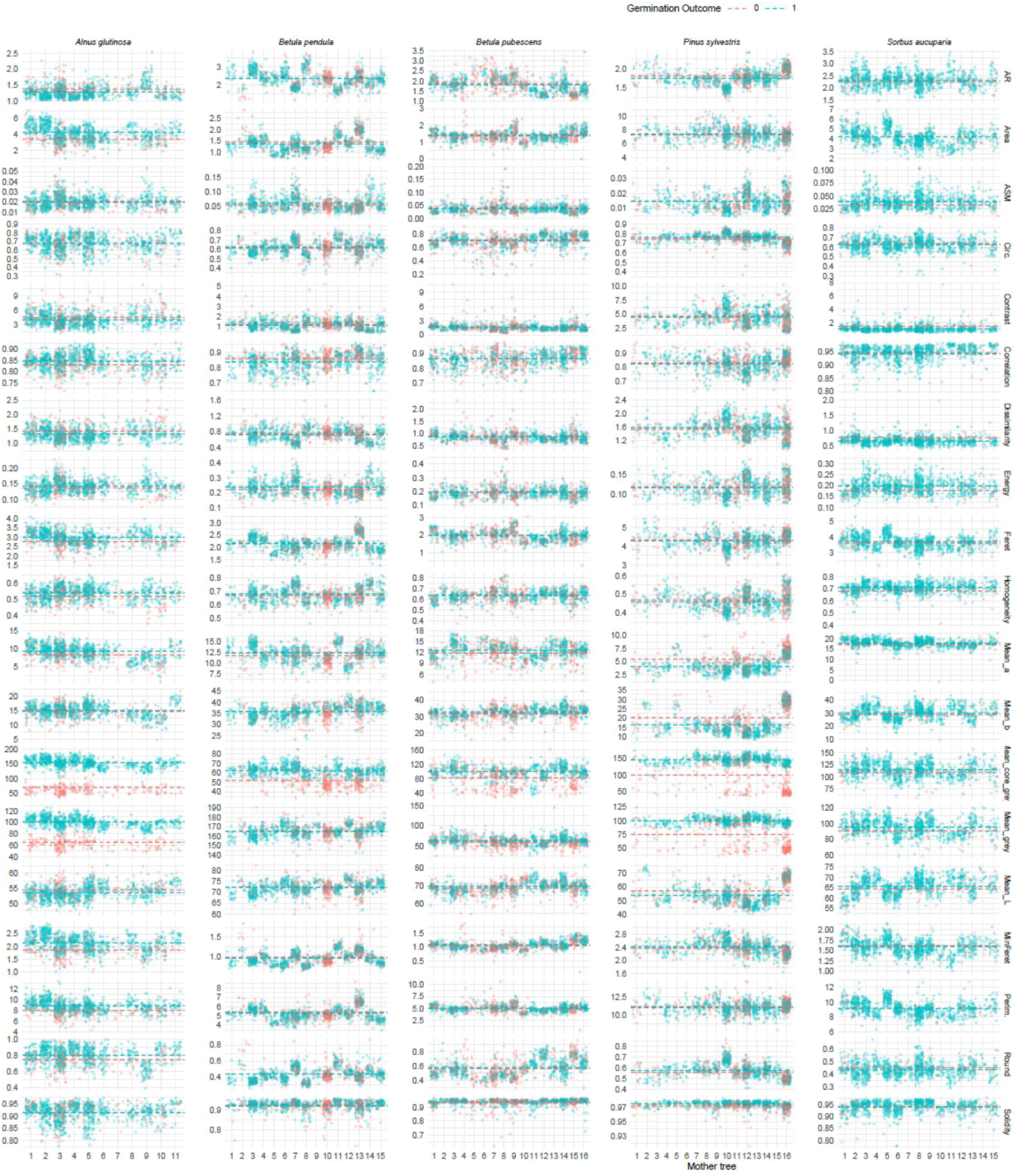
Distribution of seed morphological trait values between germinated and non-germinated seeds across individual mother trees from all species.

**Figure S3.**
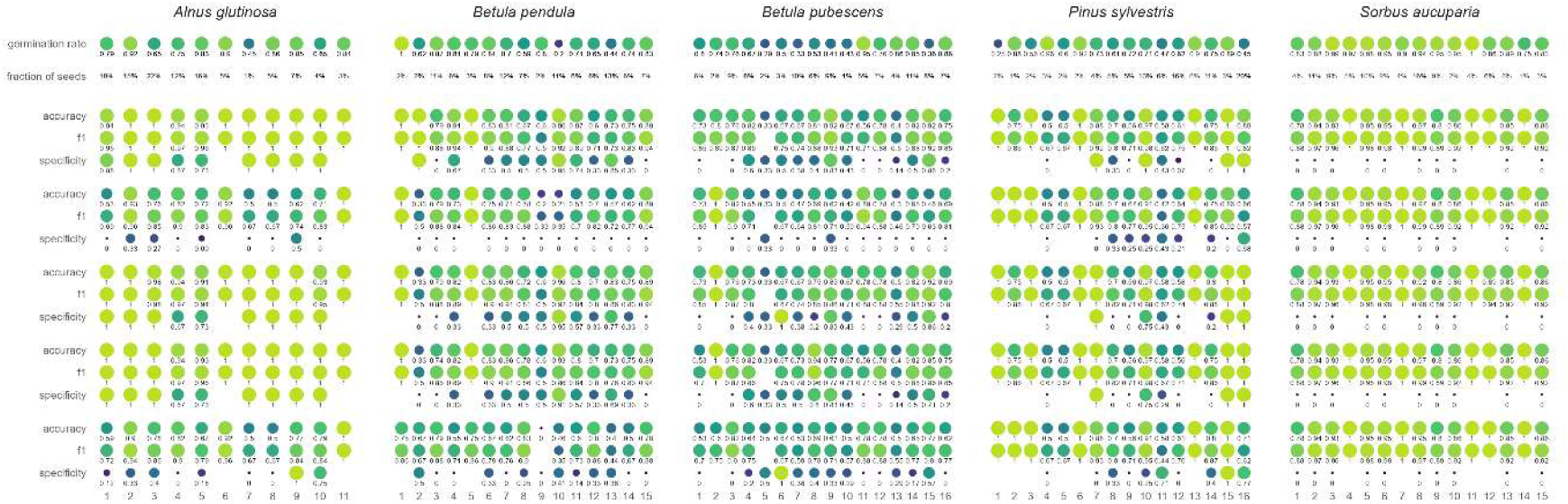
Summary of metrics obtained from models for individual mother trees of all species.

