## Supplementary figures and images for "Enhancing tree seed germination prediction with image-driven machine learning models"

### FigS1_high_resolution

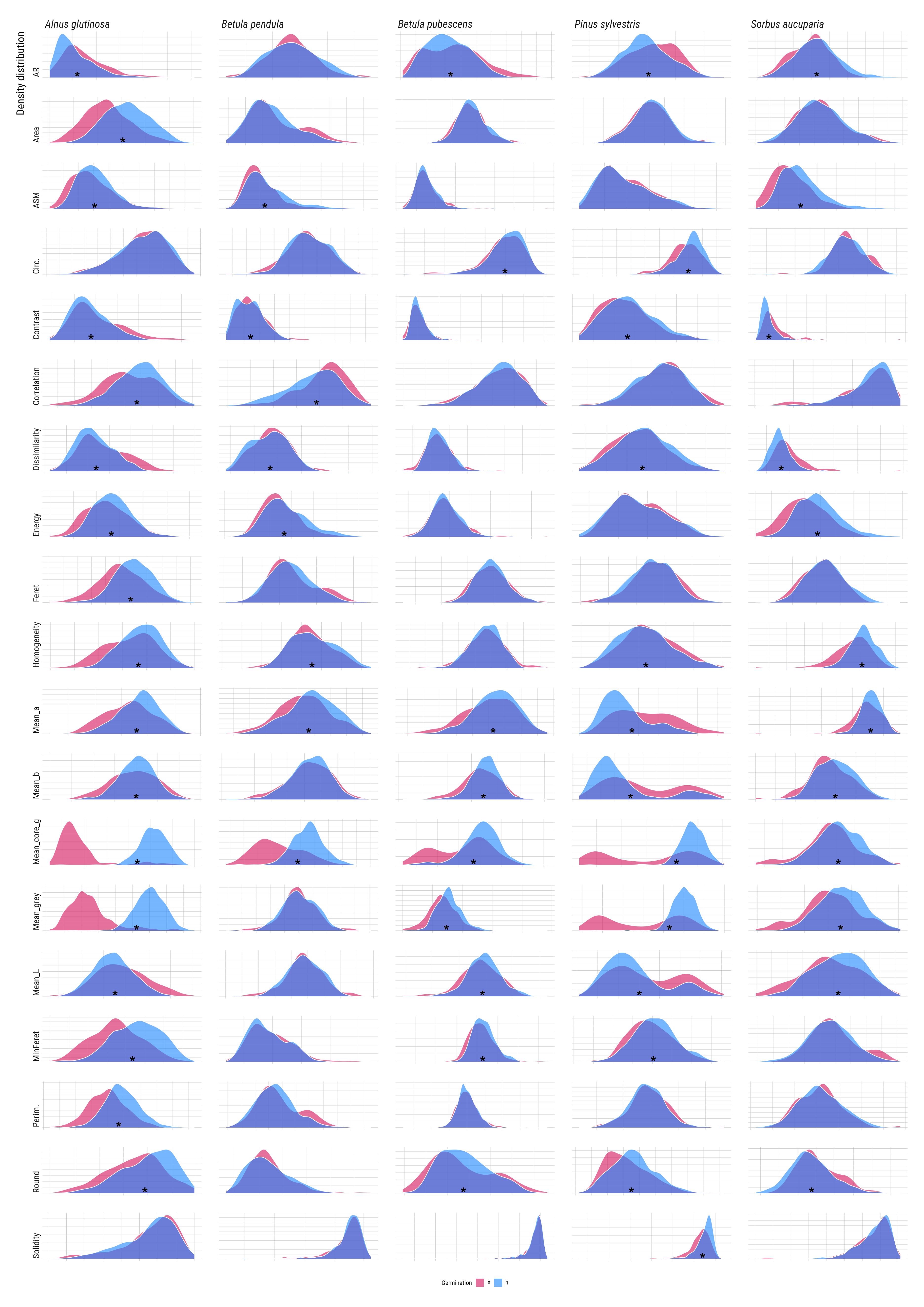

### FigS2_high_resolution

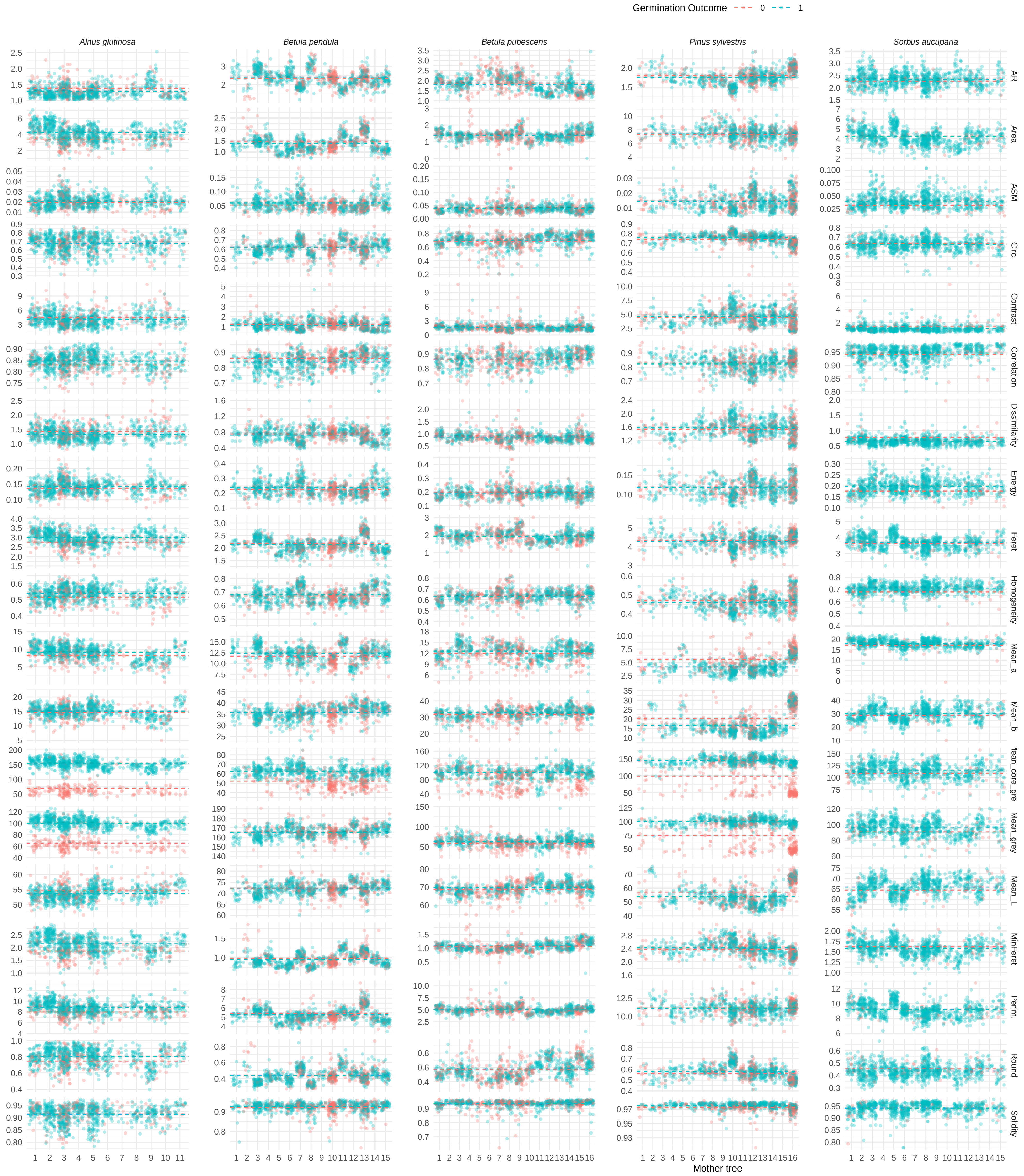

### FigS3_high_resolution

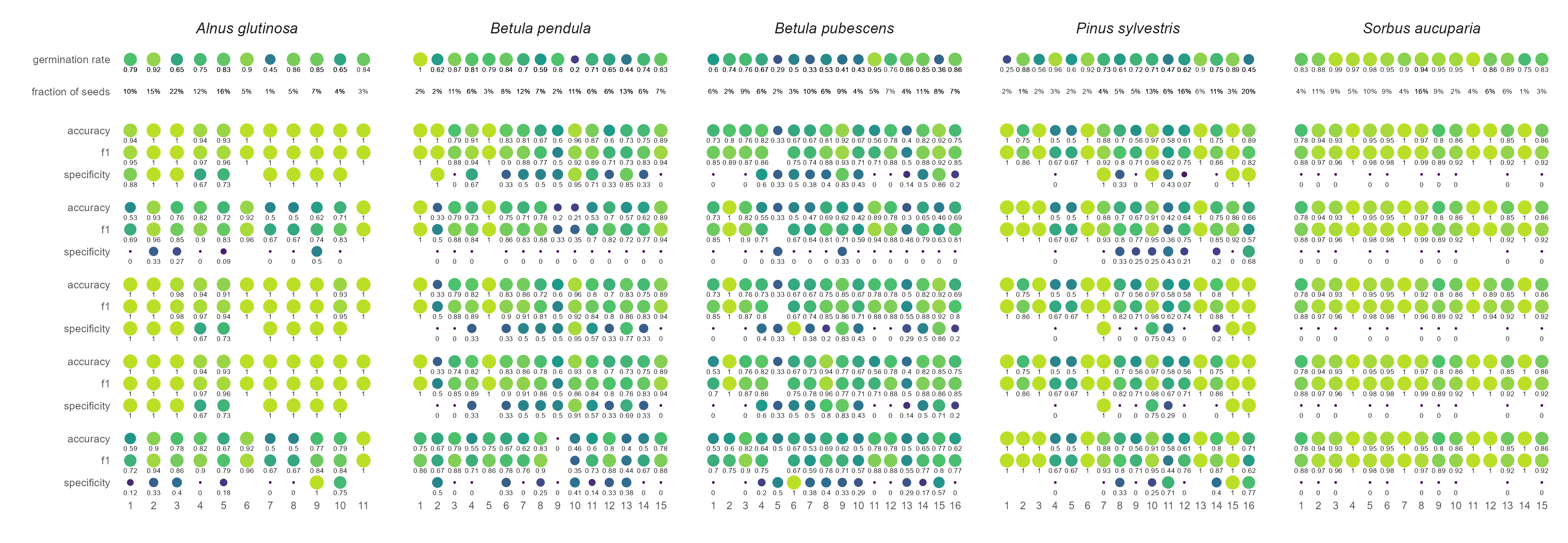
